# Single molecule co-occupancy of RNA-binding proteins with an evolved RNA deaminase

**DOI:** 10.1101/2022.09.06.506853

**Authors:** Yizhu Lin, Samentha Kwok, Bao Quoc Thai, Yewande Alabi, Megan S. Ostrowski, Ke Wu, Stephen N. Floor

## Abstract

RNA-protein interactions broadly regulate gene expression. To understand RNA regulation, it is critical to measure RNA-protein interactions in cells. Current approaches to measure RNA-protein interactions often rely on crosslinking and shortread RNA sequencing, which has considerably advanced the understanding of gene expression but also suffers from some limitations. We present REMORA (RNA Encoded Molecular Recording in Adenosines), a new strategy to measure RNA-binding events on single RNA molecules in cells. In REMORA, adenosine deamination serves as a molecular record of RNA-protein interactions that are identified by mutations by sequencing. We performed RNA-based directed evolution to identify an RNA deaminase variant with high activity on arbitrary adenosine residues in RNA. We show that this RNA deaminase has high activity, low local sequence or structure bias, low background, and is generally applicable to diverse RNA-binding proteins. By combining our improved A-to-I RNA deaminase with the C-to-U deaminase APOBEC1 and long-read RNA sequencing, our approach enables simultaneous recording of the locations two RNA binding proteins on single mRNA molecules. Orthogonal RNA molecular recording of two Pumilio family proteins, PUM1 and PUM2, reveals that PUM1 competes with PUM2 for some but not all Pumilio binding sites in cells, despite having the same *in vitro* binding preferences. Our work thus measures competition between RNA-binding proteins for RNA sites in cells, and our genetically encodable RNA deaminase enables single-molecule identification of RNA-protein interactions with cell type specificity.

## Introduction

Regulation of gene expression is critical for proper cellular function. RNA-binding proteins (RBP) play important roles in the regulation of gene expression by interacting with cellular RNA molecules across their life cycle. RBPs bind RNA to regulate mRNA splicing and polyadenylation, nuclear export, mRNA translation, and mRNA degradation^1^. Measuring the sites where RBPs bind RNA molecules in cells is necessary to understand how they regulate RNA biology. Since genes often generate multiple RNA products, it is also important to measure RBP sites with transcript isoform and singlemolecule resolution.

Several high-throughput short-read sequencing methods exist to identify RBP binding sites in RNA. RNA bind-n-seq, RNA-MaP, and related methods enable identification of RBP sequence preferences using purified proteins^2,3^. In cells, RNA immunoprecipitation (RIP-seq) and crosslinking based methods such as CLIP^4^ related methods including eCLIP^5^, PAR-CLIP^6^ identify RBP binding sites transcriptome-wide. These methods have been developed for short-read RNA sequencing and require RNase digestion of RNA molecules such that only RBP-associated regions will be purified and sequenced. However, RNase digestion limits resulting data to short fragments of RNA. Bulk-averaged short read sequencing data averages binding events across RNA molecules and results in the loss of RNA transcript isoform information. As human RNA molecules are heterogeneous and vary widely in translational potential^7,8^, RBP binding information that is averaged across RNA molecules can be challenging to interpret. Moreover, it is only possible to measure binding of one RBP at a time with these approaches, making it challenging to measure competition or cooperation between RBPs.

To distinguish RNA isoforms and measure RBP:RNA interactions at the single molecule level, recent methods including TRIBE^9^ and DART-seq^10^ or STAMP-seq^11^ have been developed. These methods use RNA deaminase enzymes as RNA molecular recorders to mark RBP binding sites. In TRIBE, the catalytic domain of the adenosine deaminase ADAR2 is fused to an RBP to introduce A-to-I editing at RBP binding sites^9^. Similarly, in DART/STAMP-seq, the C-to-U deaminase APOBEC1 is fused with a RBP of interest and introduce C-to-U mutations adjacent to RBP binding sites^10,11^. However, both ADAR and APOBEC1 have limitations as RNA molecular recorders. TRIBE relies on a drosophila ADAR enzyme, which is a double-stranded RNA deaminase that has reduced editing efficiency on single-stranded RNAs^9^. APOBEC1 has endogenous RNA binding targets especially in 3’ UTR regions of mRNAs^12^, and thus may introduce background C-to-U editing. These limitations motivate development of a synthetic RNA molecular recorder.

Here, we use RNA-based directed evolution to develop an RNA adenosine deaminase with high activity, low background, and low sequence or RNA structure preference – which we term the RNA Adenosine Base Editor (rABE). Using a stringent computational pipeline to identify induced editing sites in short- and long-read sequencing data, we validate that rABE recapitulates known binding motifs of multiple RNA-binding proteins. To facilitate description of the approach, we name this strategy REMORA: RNA Editing Molecular Recording in Adenosines. We then use rABE and APOBEC1 to measure co-occupancy and competition at the singlemolecule level between Pumilio family RNA-binding proteins that recognize the same RNA sequence. We find that the Pumilio protein with higher expression dominates binding at the population level, but singlemolecule data reveals that individual molecules can be bound by either Pumilio protein. Inducible expression also demonstrates that deamination frequency is a function of protein concentration, indicating REMORA reports on concentration-dependent binding in cells. REMORA thus enables single-molecule detection of RNA-binding proteins in cells through genetically encodable RNA adenosine deamination.

## Methods

### General molecular cloning

Gene fragments coding for tested adenosine deaminases (human *ADAR2*, human *ADAT2–ADAT3, S. cerevisiae* TAD2– TAD3, *S. aureus* tadA) were synthesized by Twist Biosciences. DNA amplification was performed with Phusion High-Fidelity PCR Master Mix (Thermo Fisher #F630) or KAPA HiFi HotStart ReadyMix (Roche) as described in manufacturer’s protocol, unless noted otherwise. PCR products were digested with 1 μl DpnI (NEB # R0176L) for 1 hour and purified by agarose gel extraction (MinElute Gel Extraction Kit, Qiagen #28606). Gibson cloning was done with NEBuilder HiFi DNA Assembly Master Mix (NEB #E2621L) or Gibson Assembly Master Mix (NEB E2611L) to insert deaminase regions into pICE backbone (Addgene #46960). Assembled plasmids were transformed into Mach1 competent cells (Invitrogen #C862003) for amplification and purified with miniprep (Qiagen) or midiprep (ZymoPURE II Plasmid Midiprep Kit). Plasmids encoding ABE7.10 (Addgene #102919) and hyperTRIBE (Addgene # 154786) were obtained from Addgene. The GFP-UAG-BoxB-mCherry was generated by inserting the UAG-BoxB cassette into pcDNA5-EGFP-NLS-P2AT2A-mCherry-PTS1 (Addgene #87828) with around-the-horn PCR^13^ as previously described.

### General mammalian cell culture

HEK 293T cells were cultured in DMEM with 4.5 g / L glucose, L-glutamine & sodium pyruvate (Corning #10-0130CV) and 10% FBS (Avantor #97068-085) at 37 °C, 5% CO_2_. Cell passage was done with trypsin (Corning #25-053-CI) with 2-5 min digestion at room temperature.

### Flow cytometry measurement of editing efficiency

Plasmids containing the GFP-UAG-BoxB-mCherry reporter and the putative deaminase were co-transfected into HEK 293T cells. Cells were seeded in 12 or 24 well plates at day 0 and reached 70-90% confluency at day 1 for transfection. Transfection was done with lipofectamine (Invitrogen) following manufacturer’s protocol. Cells were collected after 48 hours, washed with 1xPBS, and resuspended in 1xPBS containing 0.2 μg mL^-1^ DAPI to mark dead cells. Flow cytometry was performed using Fortessa SORP to measure the florescent signal of GFP (488nm 50mW laser, Blue B detector), mCherry (561nm. 50mW laser, YG D detector) and DAPI (405nm 100mW laser, Violet F detector). Positive cell populations were gated with wild type HEK 293T cells as negative controls and mono-color HEK 293T cells expressing only GFP or mCherry, after filtering for single cells. A-to-I editing efficiency was measured as the percentage of GFP+mCherry+ population in GFP+ population, and as the ratio of the mCherry to GFP signal in the GFP+ population.

### Directed protein evolution

To determine the optimal kanamycin concentration for selection, we generated pET15b-UAG-BoxB-KanR-ABE7.10 plasmid and pET15b-UAG-BoxB-KanR-ABE7.10-E59A plasmid, which encodes an inactive mutant of ABE7.10 as a negative control. We also generated a pET15b-UGG-BoxB-KanR-ABE7.10 plasmid that allows constant expression of KanR as positive control. Each of these plasmids were transformed into Mach1 competent cells, cultured in LB medium with 0, 0.3, 0.375, 0.5, and 0.75x of normal concentration of kanamycin (50 mg/L), 37 °C. Cell growth was measured as OD600 over 48 hours incubation time. 0.375x kanamycin concentration was the highest concentration that allowed for pET15b-UAG-BoxB-KanR-ABE7.10 cell growth and completely inhibited the growth of pET15b-UAG-BoxB-KanR-ABE7.10-E59A negative control cells.

The ABE7.10 mutant library was generated with error prone PCR using GeneMorph II (Agilent #200550) in the high mutation rate condition as described in manufacturer’s protocol. For each round of mutagenesis PCR, eight 25 μl reactions were performed, purified by agarose gel extraction (MinElute Gel Extraction Kit, Qiagen #28606) and pooled together. The ABE 7.10 mutant library was inserted into pET15b-UAG-BoxB-KanR plasmid downstream of lac promoter by large scale Gibson reaction and transformation. Ninety-six 12 μl Gibson reactions were performed in 96-well PCR plates using Gibson Assembly Master Mix (NEB E2611L). The Gibson reaction mix was transformed into Mach1 competent cells (4 μl Gibson into 40 μl cells, 288 reactions in 3×96-well plates) following manufacture’s protocol (Invitrogen C862003). After transformation, cells were incubated in 100 ml LB medium (US Biological, #L1505) with 100ug/ml ampicillin at 37 °C overnight, and plasmids were purified with Midiprep (ZymoPURE II Plasmid Midiprep Kit). A small volume of transformed cells (40 μl from 5 ml recover medium) was diluted 1:100, 1:1000 and 1:10000 and plated on Amp+ LB-agarose plates to evaluate transformation efficiency, and from this we estimated we had ~1.75×10^4^ colonies in total.

For kanamycin selection experiments, plasmids containing the ABE 7.10 mutant library were transformed back into Mach1 cells in a similar strategy and we obtained ~1x×10^7^ colonies in total. Cells were cultured in 100 ml LB medium with 0.375x kanamycin (18.75 mg/L), 37 °C overnight. Enriched plasmids were purified with midiprep (ZymoPURE II Plasmid Midiprep Kit), and the ABE 7.10 mutant region was PCR amplified with Phusion High-Fidelity PCR Master Mix (Thermo Fisher) for 20 cycles and sequenced by Genewiz amplicon-EZ sequencing. Mutations enriched in first round selection were cloned into the selection plasmid individually and used as template for GeneMorph error-prone PCR for second round selection.

### Stable cell line generation

Lentiviral plasmids expressing rABE or APOBEC1 were generated by inserting the deaminase region into pSLIK-TT-RBFOX2 plasmid backbone (Addgene #59770) with Gibson cloning. Lentiviral plasmids expressing RBP-rABE or RBP-APOBEC1 were generated with similar strategy. The Rbfox2 coding region was from the pSLIK-TT-RBFOX2 plasmid (Addgene #59770), PUM1 was PCR amplified from pMAL-PUM1 (Addgene #120384) and PUM2 was PCR amplified from pMAL-PUM2 (Addgene #120385). For PUM1 plasmids, the hygromycin resistance gene in the plasmid was replaced with a puromycin resistance gene for dual selection with PUM2 plasmids. These plasmids were transfected together with lentiviral packaging plasmids (1.5 μg lentiviral plasmid, 0.1 μg pGag/Pol, 0.1 μg pREV, 0.1 μg pTAT, 0.2 μg pVSVG per 1 reaction in a 6-well plate well) into HEK 293T cells with TransIT-LT1 transfection reagent (Mirus #2304) following manufacturer’s protocol. Cell culture supernatant containing lentivirus was collected and filtered after 2 days for spinfection to transduce HEK 293T cells. Briefly, 200-400 μl viral supernatant and 8 μg ml^-1^ polybrene was added in DMEM medium to total volume of 1 ml per well in 12-well plates, cells were centrifuged at 1000xg, 30 °C for 2 hours and returned to normal cell culture condition. 2 Days after spinfection, cells were treated with either 200 ug ml^-1^ hygromycin B (Gold Bio #H-270-1) or 2 μg ml^-1^ puromycin (Gold Bio #P-600-100) for at least 5 days. For dual labeling cells, we first generated HEK 293T-Tet-PUM2-rABE or HEK 293T-Tet-PUM1-APOBEC1 cells with hygromycin selection, PUM1-rABE or PUM1-APOBEC1 were then introduced by a second round of puromycin selection. Cells were treated with 0, 50, 100, 1000 ng / mL doxycycline (abcam # ab141091) for 2 days before western blot or RNA extraction.

### Western blot

Cells were lysed with RIPA buffer (Thermo Fisher # 89900) following manufacturer’s protocol. Western blots were performed with 4-20% Mini-PROTEAN(R) TGX(tm) Precast Protein Gels (Bio-rad, #4561095 or #4561096) and Trans-Blot ^®^ Turbo™ Mini PVDF Transfer Packs (Bio-rad #1704156). After membrane transfer, PVDF membranes were blocked with 5% milk in PBST buffer for 1 hour, incubated with primary antibody at 4 °C overnight, then incubated with secondary antibody at room temperature for 1 hour. Antibodies and concentration used in this study are: Monoclonal ANTI-FLAG® M2-Peroxidase (HRP) (Sigma-Aldrich #A8592) at 1:50k; anti-RBFox2 (abcam ab57154) at 1:500 – 1:1k; anti-PUM1 (abcam #ab3717) at 1:5k – 1:10k; anti-PUM2 (abcam #ab92390) at 1:1k; anti-β-Actin-HRP (Santa Cruz Biotech #sc-47778) at 1:5k; anti-β-Actin-680 (Santa Cruz Biotech #sc-47778-AF680) at 1:5k; IRDye Goat anti mouse 800CW (Licor #926-32210), IRDye Donkey anti rabbit 680RD (Licor #926-68073), IRDye Donkey anti goat 800CW (Licor #926-32214).

### RNA extraction and high-throughput RNA sequencing

Cellular RNA extraction was performed with Direct-zol RNA miniprep Kit (Zymo Research #R2053) following manufacturer’s protocol. RNA quality was verified with Bioanalyzer (Agilent RNA 6000 Pico) before library prep. For Illumina high-through sequencing library preparation, we used NEBNext® rRNA Depletion Kit (Human/Mouse/Rat) (NEB #E7405), NEBNext® Ultra™ II Directional RNA Library Prep Kit for Illumina® (NEB #E7765) and NEBNext Multiplex Oligos for Illumina (NEB #E6440S and NEB #E6442S) following manufacturer’s protocol. Sequencing was performed with a NovaSeq (S4 300 flow cell, paired-end 2×150bp) at the Center For Advanced Technology at the University of California, San Francisco.

### Long-read sequencing

Pac-bio long-read sequencing library was done with the SMRTbell Express Template Prep Kit 2.0 (Pacbio) with the Iso-seq protocol as described by manufacturer. RNA samples used in long-read sequencing were the same as used in Illumina sequencing, as described above. We used a customed TSO oligo containing 8 nt UMI barcodes for de-duplication: 5’ GCAATGAAGTCGCAGGGTTG [UMI N*8] HHHHHHHHrGrGrG 3’. Sequencing was performed with a shared PacBio Sequel II system on a SMRT Cell 8M. Long-read sequencing alignment was performed with Pacbio’s IsoSeq3 pipeline (https://github.com/ylipacbio/IsoSeq3).

### High throughput sequencing data analysis

All scripts used in this study will be available. Sequencing reads were aligned with STAR^14^ to GRCh38 human genome reference and processed with samtools^15^, picard MarkDuplicates, and GATK SplitNCigarReads^16^. Differential analysis was performed with Rsubreads and edgeR^17^. Variant calling was performed with bcftools^18^:

~~~
bcftools mpileup -f [input_reference.fa] -R
[input_annotation.bed] -d 10000000 -I -a
DP,AD,ADF,ADR,SP,INFO/AD,INFO/ADF,INFO/ADR
[input.bam]| bcftools filter -i ‘INFO/AD[1-]>2
 & MAX(FORMAT/DP)>50 -O v - > [output.vcf]
~~~

Variants were annotated with VariantAnnotation^19^ (R Bioconductor).

Significant editing site calling was done on three replicates of treated cells and three replicates of control cells using the Cochran–Mantel–Haenszel test using allele read counts and total read counts at each nucleotide location similar to previously described^20,21^. P-values were adjusted with the Benjamini-Hochberg method.

Sequence motif analysis around editing sites was performed with seqLogo (R, Bioconductor) in +/− 5nt region. De novo sequence motif discovery was done with Homer^22^ in +/− 50 nt region around editing sites. CentriMo^23^ analysis was performed with the MEME Suite (https://meme-suite.org/meme/tools/centrimo).

RNA secondary structure modeling was done with RNAfold^24^ in +/− 50 nt region around editing sites with default parameters. Random background sequences were generated by randomly sampling 2000 sequences (100 nt in length) from human cDNA sequences (Ensembl 107, GRCh38).

GSEA analysis was performed with clusterProfiler^25^ (R, Bioconductor):

~~~
gseGO(geneList = geneList, OrgDb= org.Hs.eg.db,
ont = “BP”, keyType = ‘SYMBOL’, minGSSize =
5, maxGSSize = 800, pvalueCutoff = 0.05,
pAdjustMethod = “fdr”)
~~~

The input geneList was generated by ranking genes with mean editing rate changes in different conditions.

## Results

### Identifying a single-stranded RNA adenosine deaminase

We first surveyed known RNA adenosine deaminase families for enzymes that might function on singlestranded RNA (ssRNA). We selected variants of five families of A-to-I deaminases: the human tRNA deaminase *ADAT2–ADAT3*, the *S. cerevisiae* tRNA deaminase TAD2–TAD3, the prokaryotic tRNA deaminase *S. aureus* tadA, the ADAR2-based engineered RNA deaminase hyperTRIBE^26^, and an engineered single-stranded DNA A-to-I deaminase ABE7.10^27^, which was developed using directed protein evolution from *E. coli tadA* for DNA A-to-I editing as a dCas9 fusion. The selected tRNA deaminases target a loop region of tRNA in a single stranded conformation, while ADAR2 targets double-stranded RNA (dsRNA). We therefore also included structurally-guided mutants of human ADAR2 in an attempt to retain deamination activity while reducing dsRNA preference (not shown).

To evaluate the ability of selected RNA deaminases to target ssRNA regions, we designed a fluorescent A-to-I editing reporter as shown in Figure 1A, inspired by prior work^28^. The deaminases were fused to a lambdaN peptide that interacts with the BoxB RNA element and recruits the deaminase to the adjacent UAG stop codon between eGFP and mCherry. The UAG stop codon is predicted by RNAfold to be in a single-stranded RNA conformation (Supplementary Figure 1A). When deaminated, the UAG stop codon is modified to an UIG codon, which is decoded as UGG (tryptophan), allowing readthrough of a translating ribosome to express mCherry. We transfected six putative ssRNA deaminases into HEK 293T cells along with a plasmid expressing a eGFP-UAG-BoxB-mCherry mRNA reporter. The A-to-I editing efficiency was measured using flow cytometry by the percentage of mCherry+ population in the GFP+ population. As shown in Figure 1B and 1C, all six deaminases tested showed more mCherry-positive cells compared to the empty vector control, suggesting some level of A-to-I editing. The ssDNA deaminase ABE7.10 showed the highest mCherry-positive percentage (71.6% ± 2.8%; Student’s t, α=0.05), significantly higher than other deaminases including hADAR2 (45.1% ± 1.37%), hyperTRIBE (42.2% ± 5.97 %) and *S. cerevisiae* TAD2–TAD3 (46.8% ± 3.93%). This is consistent with previous reports of the RNA editing activity of ABE7.10 when attached to dCas9^29–31^, but also indicates its deamination can be directed to specific sites. We tested the distance dependence for deamination by varying the distance between the UAG stop codon and BoxB stem-loop and found that the optimal distance was around 11nt (not shown). We then directly tested the editing efficiency of ABE7.10 using Sanger sequencing (Figure 1D) and RNA-seq (Figure 1E). We found that ABE7.10 had a maximum editing efficiency by RNA-seq of 25.8% (std 7.5%) at 11 nucleotides upstream of the BoxB structural element. We concluded ABE7.10 is a good candidate ssRNA A-to-I editor.

**Figure 1:**
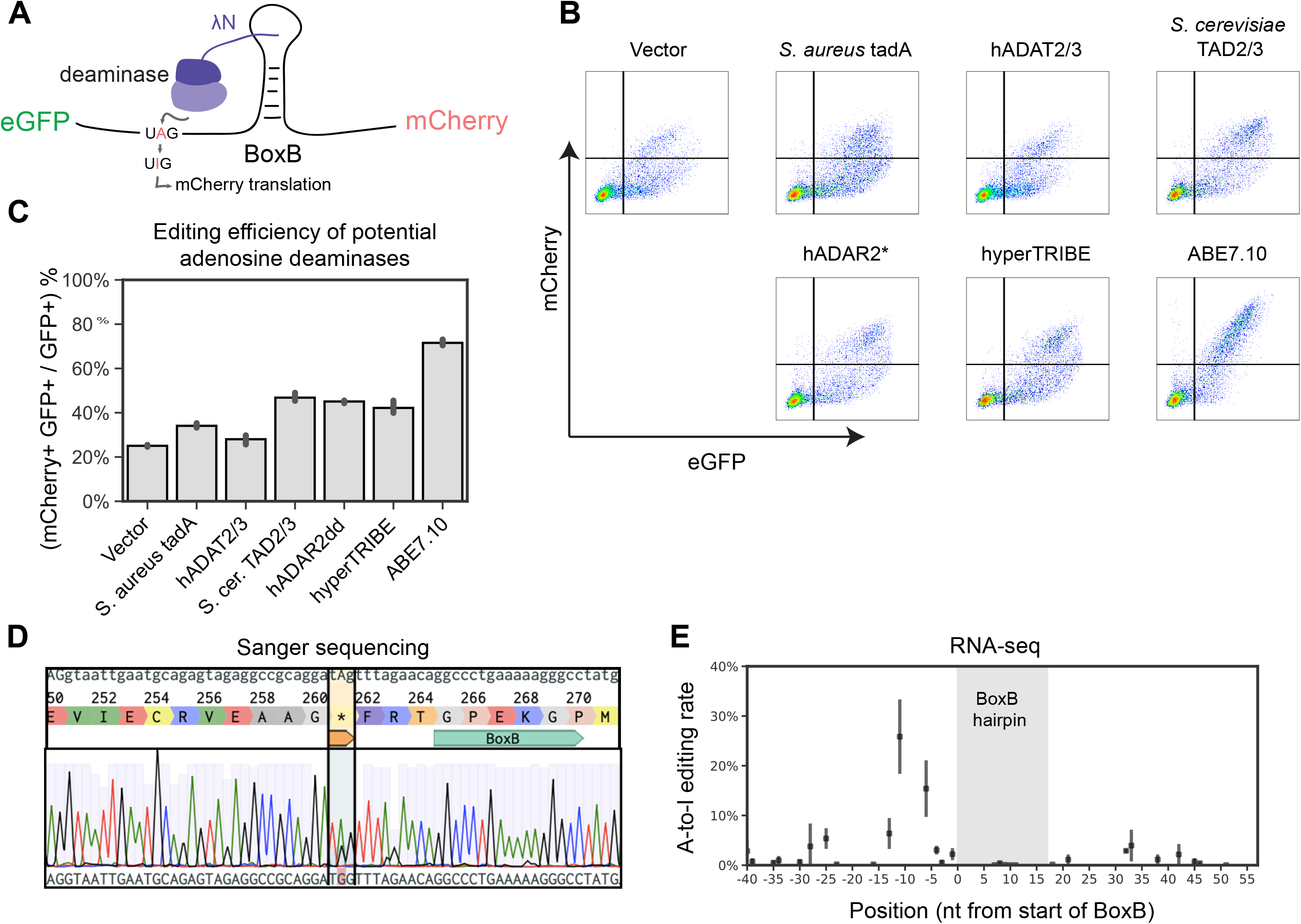
Screening for a single-stranded RNA adenosine deaminase. (A) Design of the eGFP-UAG-BoxB-mCherry deamination reporter system. Putative ssRNA deaminases were fused to a lambdaN peptide to facilitate recruitment near a stop codon by binding to a BoxB stem-loop structure in the eGFP-UAG-BoxB-mCherry reporter mRNA. Adenosine deamination at the UAG stop-codon produces UIG (where I is Inosine), which is decoded as UGG (Trp) and increases expression of mCherry. (B) Flow cytometry measurements of HEK 293T cells co-transfected with the eGFP-UAG-BoxB-mCherry reporter and six putative RNA deaminases fused to lambdaN. (C) Quantification of flow cytometry measurements by dividing the number of double-positive mCherry+ GFP+ cells by GFP+ cells as indicated by gates shown in (B). ABE7.10 shows the highest editing efficiency among tested adenosine deaminases. S. cer: Saccharomyces cerevisiae. hADAR2dd: the deaminase domain of human ADAR2. (D) Sanger sequencing identified A-to-I editing at the targeted UAG stop-codon. Note that Inosine in RNA is converted to guanosine in DNA during reverse transcription (black trace). (E) RNA-seq based measurement of A-to-I editing rates of ABE7.10 on the eGFP-UAG-BoxB-mCherry reporter mRNA after co-transfection in HEK 293T cells.

### Engineering an improved RNA molecular recorder using directed protein evolution

Since ABE7.10 was optimized for ssDNA deamination activity through directed evolution but started as a tRNA deaminase, we reasoned that it could be further mutated to increase its editing efficiency on ssRNA. We designed a directed protein evolution system to select for variants of ABE7.10 that might be an improved ssRNA A-to-I editor. A UAG-BoxB cassette was inserted at the 5’ end of the kanamycin resistance gene *kanR* (Figure 2A). Similar to the eGFP-UAG-BoxB-mCherry reporter mRNA, deamination at the UAG stop codon will enable expression of *kanR*. Since edited RNA molecules are transient, this selection proved sensitive to the initial starting optical density and kanamycin concentration. To determine the optimal kanamycin concentration for RNA deaminase selection, we incubated *E.coli* expressing ABE7.10 and the inactive mutant ABE7.10-E59A with different concentrations of kanamycin in deep 96-well plates for 48 hours. As shown in Supplementary Figure 2A, 18.75 μg/ml kanamycin (37.5% of the normal working concentration) allowed for growth of ABE7.10-expressing *E.coli* and could effectively inhibit growth of inactive ABE7.10-E59A expressing *E. coli*.

**Figure 2:**
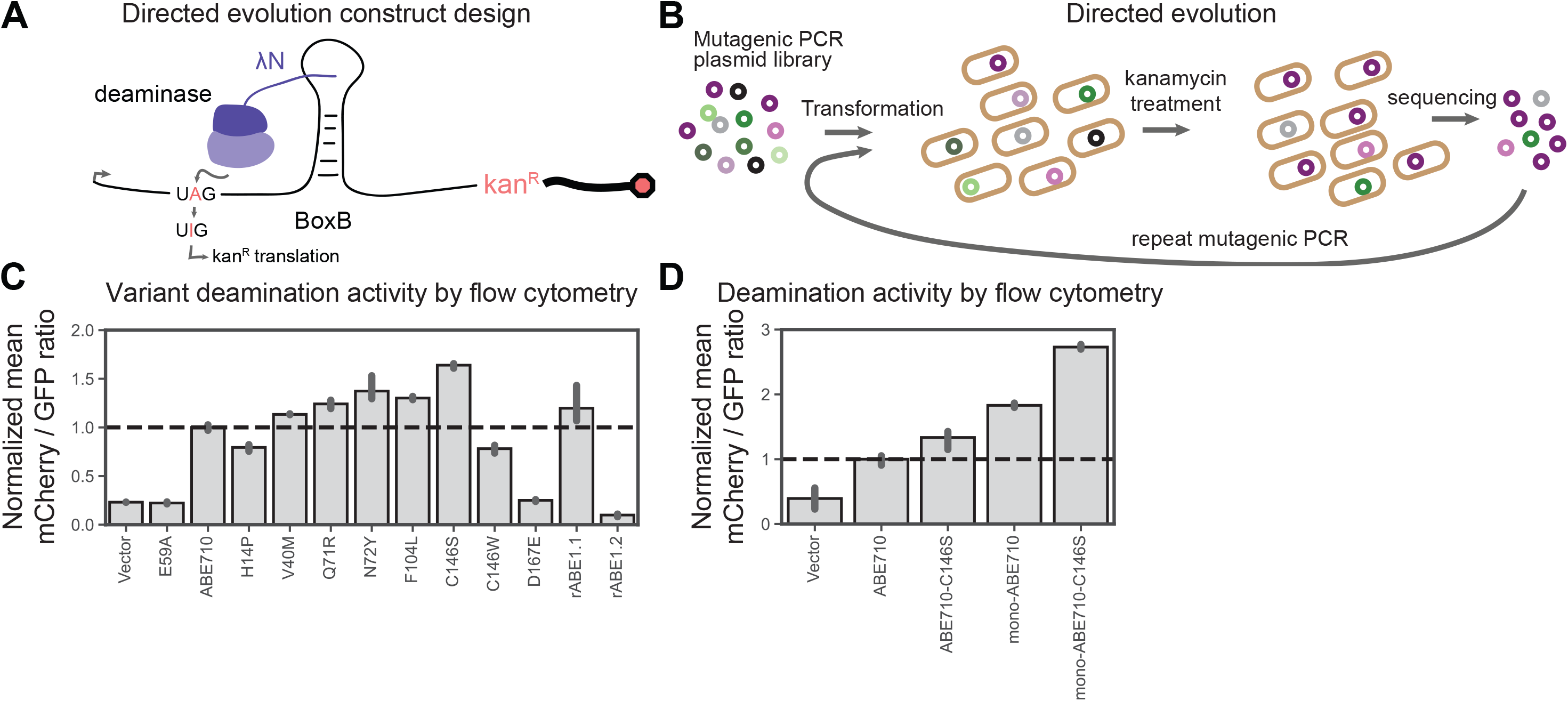
Engineering an improved ssRNA adenosine deaminase by directed protein evolution. (A) Design of UAG-BoxB-kanR reporter for directed protein evolution in E. coli. (B) Directed protein evolution workflow. Random mutations were introduced into the ABE7.10 protein coding region by error prone PCR and were transformed into E. coli Mach1 cells together with UAG-BoxB-KanR reporter. Cells were incubated with kanamycin overnight to select for ABE7.10 mutants that survived selection. Enriched mutants after selection were identified by amplicon sequencing, transformed back into E. coli cells for a second round of directed protein evolution. (C) Editing efficiency of enriched ABE7.10 mutants measured by flow cytometry with the GFP-UAG-BoxB-mCherry reporter. (rABE1.1: ABE7.10 + V40M, Q71R, N72Y, F104L, C146S; rABE1.2: ABE7.10 + V40M, R47S, G66V, Q71R, N72Y, F104L, R107L, C146S). (D) A comparison of monomeric and dimeric ABE7.10 and rABE variants using flow cytometry and the GFP-UAG-BoxB-mCherry reporter.

To search for ABE7.10 variants that might improve deamination efficiency, we introduced random mutations into the ABE7.10 coding region with GeneMorph II error prone PCR using the high mutation rate protocol (9-16 mutations/kb). We generated a library of prokaryote-expressing plasmids harboring both the BoxB-UAG-*kanR* selecting gene and randomly mutated ABE7.10 variants, transformed it into *E.coli* by large scale electroporation (megaX), and incubated the transformants in kanamycin-containing LB medium for up to 48 hours (Figure 2B). After 1 cycle of directed protein evolution, the ABE7.10 mutant region was sequenced with amplicon sequencing to identify enriched mutations. The enriched mutations are summarized in Supplementary File 1. We then cloned promising individual mutations and tested their A-to-I editing efficiency with the mGFP-UAG-BoxB-mCherry reporter and flow cytometry. Variants with higher editing efficiency than ABE7.10 were pooled for a second round of error prone PCR and selection. Initially, we used a threshold in mCherry levels to detect any editing activity. To more directly test editing level as opposed to any editing activity, we calculated the mean of ratios of mCherry to GFP fluorescence intensity in the GFP-positive population. This measure showed good sensitivity and reproducibility when measuring A-to-I editing efficiency (Figure 2C and Supplementary Figure 2B). We found that the C146S mutation increased the A-to-I editing efficiency of ABE7.10 on ssRNA by more than 1.6 fold. Combining multiple single amino acid mutations that increased editing efficiency (rABE1.1 and rABE1.2) did not outperform ABE7.10^C146S^.

Interestingly, the single mutation C146S mutates a cysteine back to serine as in wild type *E. coli* tadA. Structural modeling using *S. aureus* tadA (PDB: 2B3J)^32^ shows that Thr145 (corresponding to *E. coli* Ser146) is about 4.5 angstroms from the phosphate backbone of bound RNA, suggesting that amino acid substitutions at this position may influence RNA–protein interactions (Supplementary Figure 2C). We also observed *S. aureus* Thr145 is in the vicinity of U33, the −1 nucleotide prior to the adenosine to be deaminated. To explore the possibility that C146S influences local sequence preferences, we analyzed Sanger sequencing traces on an adenosine-rich version of the BoxB deamination reporter with numerous adenosines mutated adjacent to the target stop codon. We find that C146S increases deamination at adenosines with a −1 uracil and in other sequence contexts (such as CAG or GAA), suggesting that C146S does not induce a strong dependence for a −1 uracil (Supplementary Figure 2D).

The tRNA deaminase tadA naturally functions as a homodimer^32^. To mimic this, ABE7.10 was designed as a constitutive dimer of two tadA domains^27^. However, recent studies further optimized the DNA editing efficiency of ABE7.10 and found that a single tadA domain is sufficient to achieve high editing efficiency^33^. We therefore tested whether mono-ABE7.10^C146S^, with just a single tadA monomer, can achieve similar or higher RNA editing efficiency to dimeric tadA-ABE7.10^C146S^. Flow cytometry using the eGFP-UAG-BoxB-mCherry reporter showed that mono-ABE7.10^C146S^ has 2.1 (±0.5) fold higher A-to-I editing efficiency on ssRNA than the tadA-ABE7.10^C146S^ dimer and 2.73 (±0.07) fold of the original ABE7.10 (Figure 2D). We named this construct the RNA Adenosine Base Editor (rABE).

### rABE has low background and minimal sequence dependence

To assess background edits and the impact of rABE on gene expression, we generated HEK 293T cell lines expressing rABE under a doxycycline inducible promoter. Cells were treated with 0, 50, or 1000 ng ml^-1^ doxycycline for 48 hours before RNA extraction and RNA-seq library preparation (Figure 3A). Even under the high doxycycline condition, expression of rABE alone had no significant effect on cellular gene expression (Figure 3B; odds ratio threshold = 2, p < 0.05, BH adjusted). Therefore, induction of rABE does not induce detectable gene expression changes.

**Figure 3:**
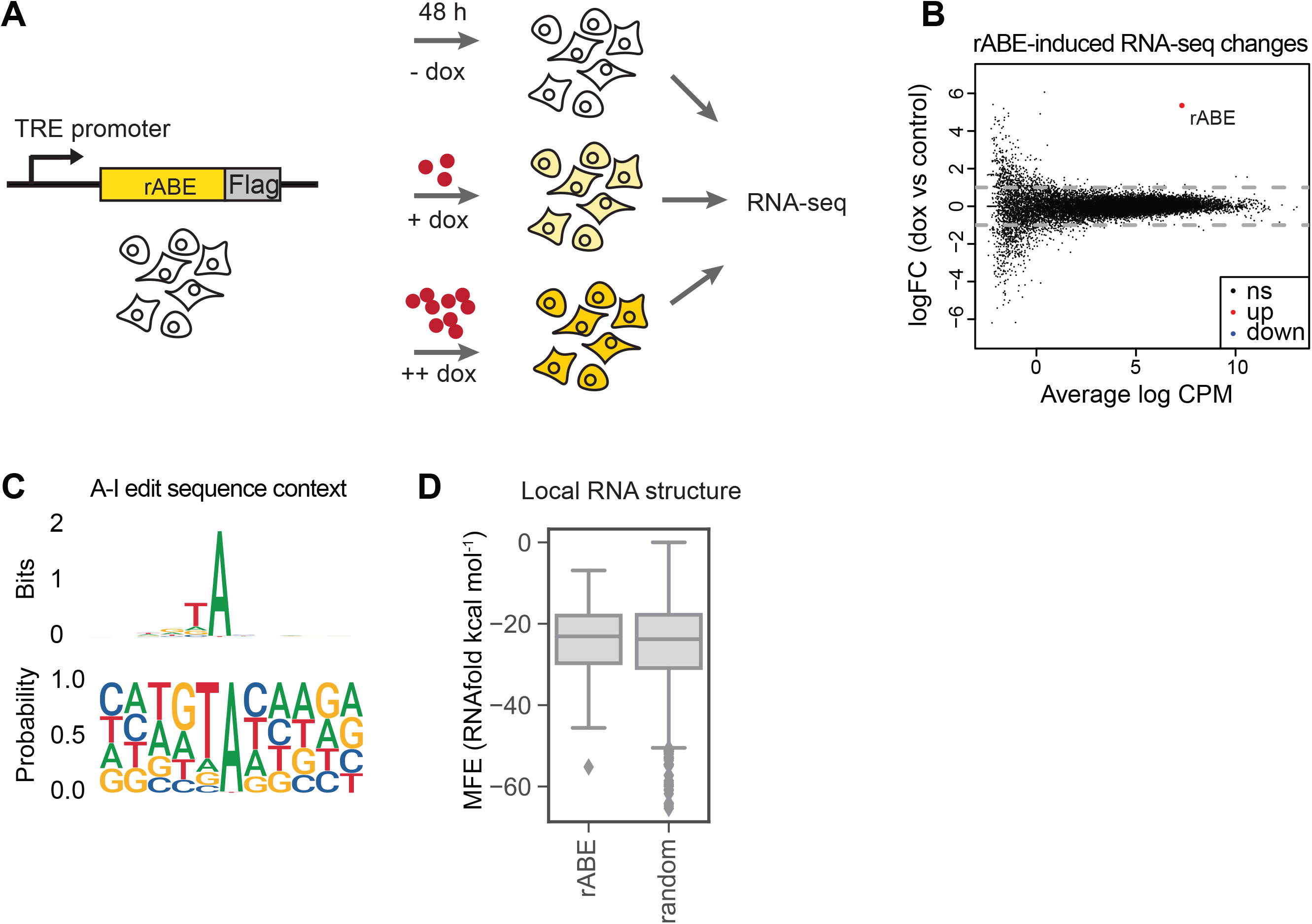
Transcriptome-wide characterization of rABE. (A) A stable HEK293T cell line expressing rABE-Flag was generated by lentiviral transduction. The expression of rABE-Flag was under regulation of Tet expression system and was induced with addition of doxycycline. Cells were incubated with 0, 50, or 1000 ng/ml doxycycline for 48 hours before RNA extraction and RNA-seq. (B) Differential expression analysis comparing transcriptome-wide gene expression level after doxycycline induction (1000 ng / ml vs no doxycycline control). Expression of rABE alone does not change gene expression levels. (C) Sequence composition averaged across A-to-I editing sites introduced by rABE. (D) MFE of predicted RNA structure in a window of 50 nt around rABE editing sites versus 2000 random background sequences.

To measure rABE editing sites, we developed a stringent computational pipeline to call A-to-I edits in RNA-seq data based on significantly increased editing over background in all three replicates. Briefly, we first filtered for read coverage (average count per sample ≥ 20) and removed SNPs or naturally-occurring A-to-I sites (allele count ≥ 2 in no dox control samples). We then performed a Cochran–Mantel–Haenszel (CMH) test on the total read count and allele count at each nucleotide position. For every significant editing event (p < 0.05, BH adjusted), the editing strength was calculated as the average of the difference in editing rate between treated and control samples.

When not fused to an RNA–binding protein, we found that expression of rABE alone introduced sparse background editing even under high doxycycline induction. We identified 365 and 247 background editing events when comparing the no doxycycline condition to the high and low doxycycline conditions, respectively. To evaluate whether rABE editing has bias towards local sequence around the deaminated adenosine, we calculated the sequence frequency around 365 identified A-to-I editing sites in the high doxycycline condition. We found that rABE favored uridine at the −1 position (68.8% frequency), where A (12.3%), G (12.1%) and C (6.8%) are less represented (Figure 3C). Despite this moderate preference, all four bases were present near editing sites, suggesting rABE was tolerant to diverse sequence compositions. To test whether rABE has RNA structure dependence, we predicted RNA secondary structure in a window of 50 nt around the 365 identified A-to-I sites. Compared to 2000 random sequences sampled from the human transcriptome, we observed no differences in the minimum free energy distribution (Figure 3D), suggesting rABE had no significant bias towards RNA secondary structure distinct from background structure. These results showed that rABE had low background editing and minimal sequence and structure dependence. To rigorously identify specific RBP-associated A-to-I sites versus background sites, we used rABE-expressing cells with the corresponding doxycycline induction as negative controls for all following RBP-rABE RNA molecular recording experiments.

### rABE recovers known binding patterns of RNA-binding proteins

To test whether rABE is a faithful RNA molecular recorder, we decided to fuse rABE to the C-terminus of RNA-binding proteins with known specificity. We fused rABE to RNA-binding proteins under the control of a tetracycline inducible promoter, induced expression with doxycycline, extracted RNA, performed RNA-seq, and used our computational pipeline to call A-to-I edits above background (Figure 4A). We decided to start with the Pumilio family proteins PUM1 and PUM2, which bind to Pumilio binding elements (PBEs) specified by a UGUANA motif^34,35^. We therefore generated separate stable HEK 293T cell lines expressing PUM1-rABE and PUM2-rABE driven by a tetracycline inducible promoter, and treated cells with 0, 50, or 1000 ng ml^-1^ doxycycline for 48 hours before RNA extraction and RNA-seq library preparation.

**Figure 4:**
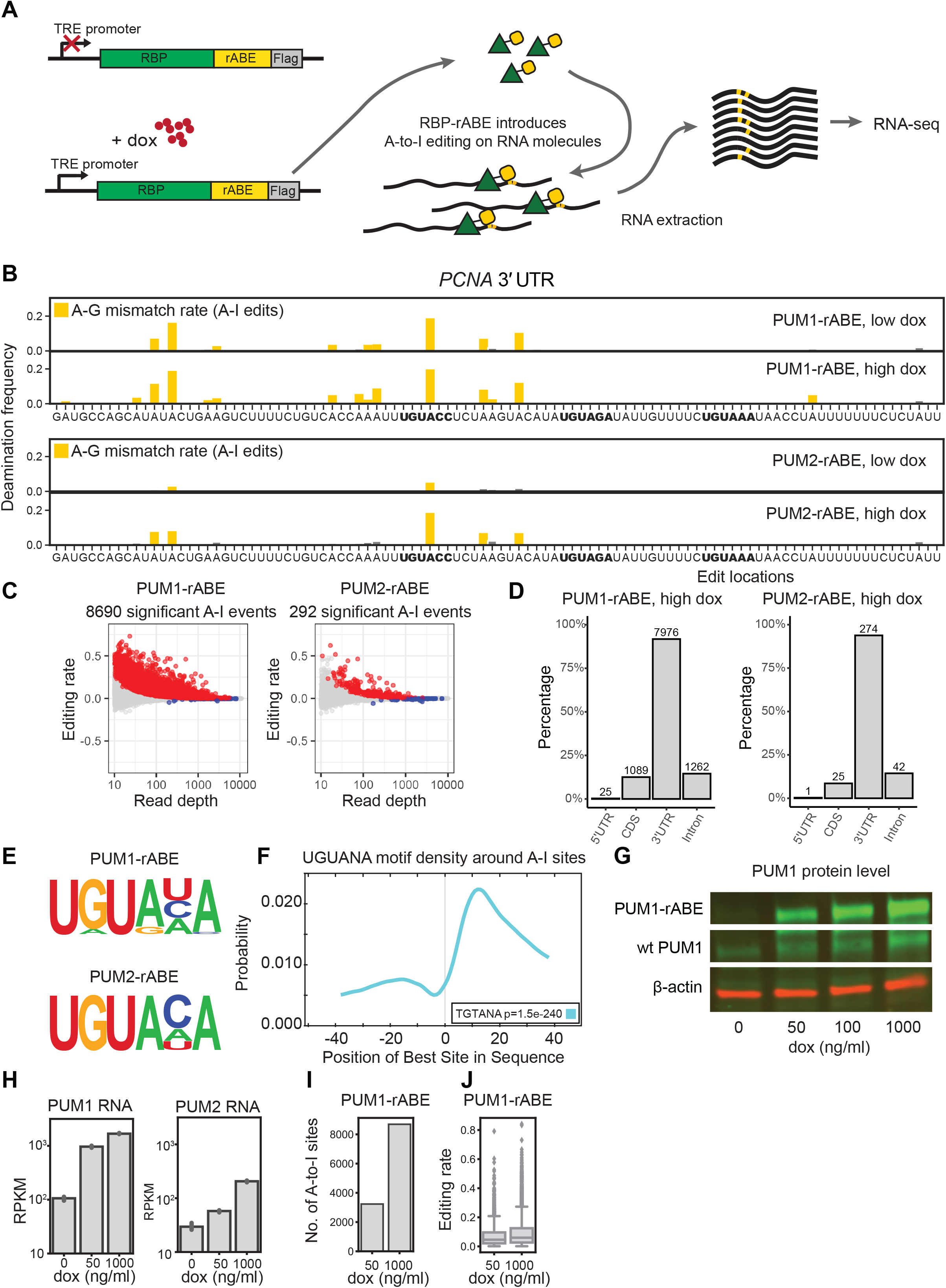
rABE is an RNA molecular recorder. (A) Stable HEK293T cells lines expressing PUM1-rABE or PUM2-rABE under a doxycycline inducible promoter were generated by lentiviral transduction. Cells were incubated with 0 (control), 50 (low), or 1000 (high) ng ml-1 doxycycline before RNA extraction and RNA-seq. PUM1-rABE or PUM2-rABE introduces A-to-I editing events adjacent to PUM1/2 binding sites that are identified as A-to-G mutations by high throughput sequencing. (B) A-to-I editing events in the 3’ UTR region of PCNA introduced by PUM1/2-rABE. The deamination frequency is the frequency of A-G transitions in PUM1/2-rABE at the indicated dox induction level over background A-G transitions in rABE alone at the corresponding dox level; A-G transitions result from A-to-I editing. Bar plots of deamination frequency are the average frequency across three replicates. Pum binding elements (PBE) motifs are highlighted in bold. (C) Differential A-to-I editing rates from PUM1/2-rABE compared to rABE-Flag untagged control under 1000 ng / ml doxycycline induction. Significant A-to-I editing events (padj < 0.05) are marked in red. (D) Transcriptome-wide distribution of significant A-to-I editing events introduced by PUM1-rABE or PUM2-rABE. The majority of editing events are located in the 3’ UTR region of targeted mRNA. As some editing sites are annotated in multiple regions in different transcript isoforms, the sum of the percentage of four categories is greater than 100%. (E) Motif enrichment in the +/− 50 nt region around A-to-I editing sites identifies canonical Pumilio binding elements. (F) CentriMo analysis around PUM1-rABE A-to-I editing sites. The PRE motif was enriched at +15 nt downstream of A-to-I editing sites. (G) Expression level of endogenous PUM1 and PUM1-rABE under 0, 50, 100, 1000 ng/ ul doxycycline induction. (H) Expression level of PUM1 or PUM2 measured by RNA-seq, under 0, 50, 1000 ng/ ml doxycycline induction. Both endogenous PUM2 and PUM2-rABE showed lower expression level than PUM1s. (I,J) Higher doxycycline concentration lead to a higher number of significant A-to-I editing events (I) and higher editing rates (J).

We found that PUM1-rABE introduced A-to-I edit sites above background in cells, as exemplified by the 3’ UTR region of PCNA, a previously identified PUM1 target^36^ (Figure 4B). PUM1-rABE A-to-I editing events were adjacent to PBE motifs and previously identified CLIP-seq peaks^37^. Transcriptome-wide, we identified 8690 significant A-to-I editing events in 1682 genes under high doxycycline condition in PUM1-rABE expressing cells (Figure 4C). 1118 out of 1682 (66.5%) of PUM1 targeted genes were also identified in previous work using CLIP-seq^37^ (Supplementary Figure 3A). Consistent with previous PUM1 footprinting with CLIP-seq, the majority of PUM1 binding events were in 3’ UTRs (7976/8690, 91.8%) (Figure 4D). We performed de novo motif identification within 50 nt around A-to-I editing sites and found that the consensus PBE motif UGUANA was enriched (Figure 4E). CentriMo^23^ analysis showed that the UGUANA motifs were on average 10 ~ 15 nt downstream of A-to-I editing sites (Figure 4F), suggesting PUM1-rABE tends to introduce A-to-I editing 10 ~ 15 nt upstream of a binding site, consistent with the rABE-lambdaN construct that used in our fluorescent reporter assay (Figure 1).

The number of PUM1-rABE A-to-I editing events identified over background increases with PUM1-rABE expression level. High doxycycline treatment (1000 ng ml^-1^) increases expression of PUM1-rABE at both the RNA and protein level (Figure 4G and 4H). We identified 3233 significant A-to-I editing events under low doxycycline treatment, 37%of the high doxycycline condition (Figure 4I), and the average editing rate at each site was slightly lower (Figure 4J), consistent with fewer molecules being edited at low doxycycline levels. This suggests that rABE deamination frequency is a measure of concentration dependent RBP binding in cells.

To test the generality of rABE, we fused rABE to the C-terminus of PUM2, another Pumilio family RNA–binding protein. We again recovered the known PBE (Figure 4D), and we observed that PUM2 identified sites have a moderate preference for cytidine at the 5^th^ position compared to PUM1 (Figure 4E). The PUM2-rABE fusion identified only 292 significant A-to-I editing events even under high dox condition (Figure 4C). Consistent with previous work^37,38^, the large majority of PUM2 targets were also PUM1 targets (112 out of 126 genes) (Supplementary Figure 3B). In the example of the *PCNA* 3’ UTR (Figure 4D), we also observed PUM2-rABE editing events near PBEs. PUM2-rABE identified many fewer A-to-I events than PUM1-rABE, and fewer PUM2-rABE targeted genes than identified previously with CLIP-seq^37^ (Supplementary Figure 3C). The lower amount of PUM2 binding events may be due to lower PUM2-rABE expression level than PUM1-rABE (Figure 4H), even under the same doxycycline condition. Notably, endogenous PUM2 also has lower expression level than PUM1 (Figure 4H). Together, we conclude that the REMORA approach is a faithful RNA molecular recorder that can be used to identify RNA-binding protein sites through introduced adenosine deamination.

### Benchmarking orthogonal deaminases using Rbfox2

To benchmark rABE on other RNA–binding proteins and compare to other deaminase recorder strategies, we fused rABE to the C-terminus of Rbfox2, a well-studied RNA–binding protein involved in splicing regulation^39–41^. We generated stable HEK 293T cell lines with tetracycline-inducible expression of Rbfox2-rABE, as above for Pumilio proteins. Following high doxycycline induction, RNA extraction and RNA-seq, we identified 3243 A-to-I editing events over background introduced by Rbfox2-rABE across 402 genes (Figure 5A). Previous work used APOBEC1, a C-to-U RNA deaminase, for RNA–binding protein molecular recording^10,11^. To compare the performance of rABE and APOBEC1, we also expressed an Rbfox2-APOBEC1 fusion in the same experimental format and performed RNA-seq. Transcriptome-wide, Rbfox2-APOBEC1 identified 1142 significant C-to-U editing events under high doxycycline condition, about one-third of Rbfox2-rABE (Figure 5A).

**Figure 5:**
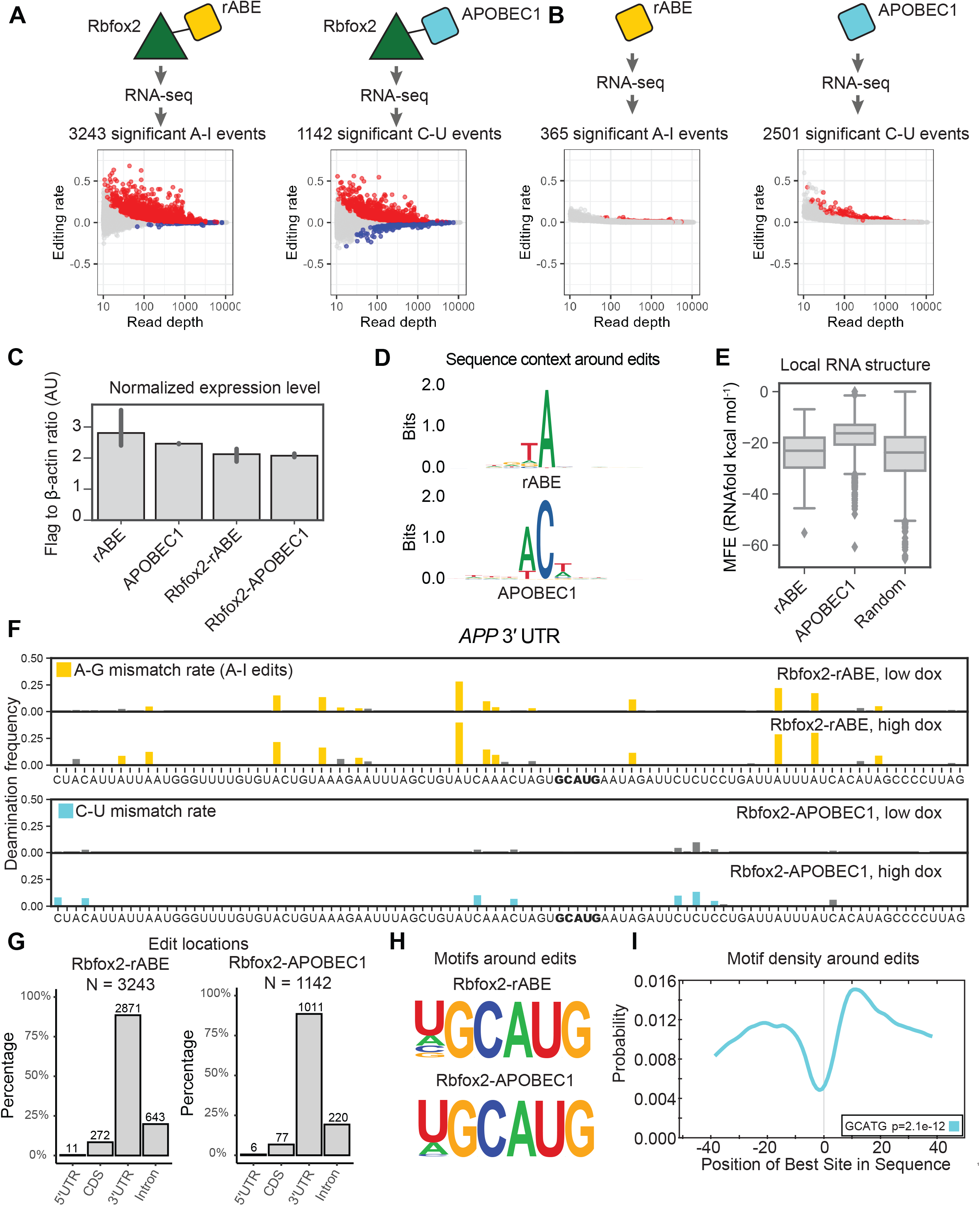
Comparison of rABE with the C-to-U RNA editor APOBEC1. (A) Transcriptome-wide Rbfox2 RNA molecular recording with Rbfox2-rABE and Rbfox2-APOBEC1 with significant editing events (padj < 0.05) in red. (B,C) Background editing from expression of rABE-Flag or APOBEC1-Flag. rABE showed lower background editing than APOBEC1 (B) at comparable expression level by RNAseq (C). (D) Sequence logos around rABE (top) and APOBEC1 (bottom) editing sites identifies local sequence preferences. (E) RNAfold structure predictions in a window of 50 nt around rABE and APOBEC1 editing sites versus 2000 random sites in the genome. (F) Example of Rbfox2-rABE and Rbfox2-APOBEC1 introduced editing patterns around the GCAUG motif in the 3’ UTR region of APP, as described in Figure 4B. (G) Distribution of Rbfox2-rABE and Rbfox2-APOBEC1 exiting sites across transcript regions. (H) Motif discovery around Rbfox2-rABE and Rbfox2-APOBEC1 recovered the known Rbfox2 binding motif, GCAUG. (I) CentriMo analysis of Rbfox2-rABE around A-to-I editing sites. Rbfox2-rABE A-to-I editing events are between 15-20 nt both upstream and downstream of Rbfox2 binding motifs.

To explore the difference in the number of identified Rbfox2 sites between rABE and APOBEC1 fusions, we measured background editing by expressing rABE or APOBEC1 alone. We explored background editing because all editing sites in this work are identified as statistically enriched over background expression of the deaminase at the same doxycycline induction level. We identified 2501 significant background C-to-U editing events introduced by APOBEC1 under high doxycycline conditions, 6.85 fold higher than background editing by rABE alone (365 A-to-I editing events; Figure 5B), despite comparable expression levels (Figure 5C and Supplementary Figure 4). We compared the local sequence preference around editing sites for rABE and APOBEC1 and find that rABE showed lower sequence bias around editing sites than APOBEC1 (Figure 5D), which prefers −1 A (83.2% frequency) and +1 U (52.0%). The local sequence preference for APOBEC1 is consistent with previous studies on endogenous APOBEC1 targets^12^. Notably, G and C are almost absent from the −1 position of APOBEC1 C-to-U editing sites, with frequency of 0.8% and 0.08% respectively, suggesting APOBEC1 has stronger local sequence dependence than rABE. RNAfold^24^-predicted structure around APOBEC1 editing sites also had higher MFE (less predicted structure) than random regions from the human transcriptome (Figure 5E, p=9.17e-263, t-test of mean), possibly suggesting some preference of APOBEC1 for unstructured RNA regions.

We identified known Rbfox2 targets in RNA editing events introduced by Rbfox2-rABE or Rbfox2-APOBEC1, as exemplified by the 3’ UTR region of *APP*, a previously identified Rbfox2 target^11,42^ (Figure 5F). Both Rbfox2-rABE and Rbfox2-APOBEC1 showed significant editing signals that were clustered in adjacent regions (Figure 5F). Transcriptome-wide, we found that 83.6% of Rbfox2-rABE and a similar percentage of Rbfox2-APOBEC1 editing sites were in 3’ UTRs (Figure 5G). This differs from prior work that used CLIP to identify Rbfox2 binding sites and found the majority of Rbfox2 sites are intronic^41,43^. One possibility for this difference is that REMORA and related approaches measure binding in the whole transcriptome, where intron-containing messages are comparatively rare, versus pull-down enrichment for RBP-associated RNAs in CLIP. Motif enrichment analysis within 100 nt regions centered around Rbfox2-rABE introduced A-to-I editing sites or Rbfox2-APOBEC1 introduced C-to-U editing sites recovered the known GCAUG motif of Rbfox2 (Figure 5H). CentriMo^23^ analysis showed that the GCAUG motif occurs at the highest frequency at 10 nucleotides downstream of A-to-I editing sites, but also has high frequency about 20 nt upstream of A-to-I editing sites (Figure 5I), possibly suggesting Rbfox2-rABE tends to introduce A-to-I editing on both sides of Rbfox2 binding sites.

### Dual color labeling with rABE and APOBEC1 for simultaneously orthogonal RBP molecular recording

Since both rABE and APOBEC1 can be used for RBP molecular recording and introduce distinguishable mutations into RNA molecules, we decided to test whether tagging rABE and APOBEC1 to two different RBPs can achieve simultaneous RBP molecular recording. We generated dox inducible stable cell lines expressing PUM1-rABE, PUM1-APOBEC1, PUM2-rABE, and PUM2-APOBEC1 individually, and cell lines that express PUM1-rABE and PUM2-APOBEC1 simultaneously. Consistent with results from Rbfox2-APOBEC1 as described above, we identified more significant editing events with PUM1-rABE (8690 sites) and PUM2-rABE (292 sites) (Figure 4C) compared to PUM1-APOBEC1 (7766 sites) and PUM2-APOBEC1 (62 sites) (Figure 6A) due to lower background editing from rABE. The choice of deaminase for RNA molecular recording had a larger impact on PUM2 than on PUM1 (89% as many sites identified in PUM1 versus 21% in PUM2). Thus, we used PUM1-APOBEC1 and PUM2-rABE RBP-deaminase combinations for the following experiments.

**Figure 6:**
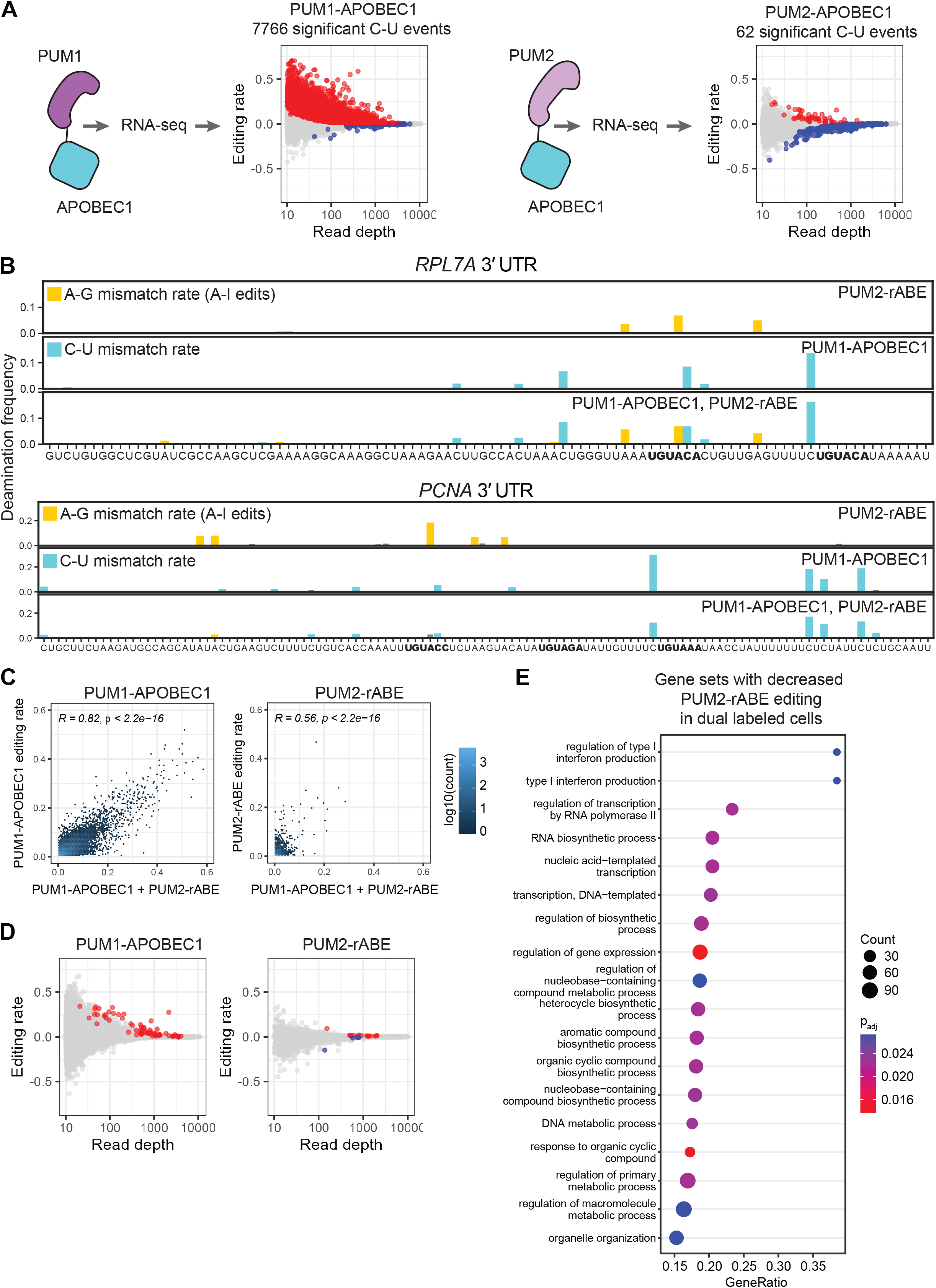
Co-occupancy and competition of PUM1 and PUM2 using rABE and APOBEC1 fusions. (A) Transcriptome-wide C-to-U editing events introduced by PUM1-APOBEC1 or PUM2-APOBEC1. Significant editing events (padj < 0.05) are marked in red. (B) Editing patterns in the 3’ UTR region of RPL7A and PCNA around Pumilio binding motifs, as described in Figure 4B. In RPL7A we observed similar editing patterns in single labeling of PUM2-rABE or PUM1-APOBEC1 and dual labeling of both, suggesting independent binding. In PCNA, we observed decreased PUM2 binding when both PUM1-APOBEC1 and PUM2-rABE are expressed, suggesting competition. (C) Scatter plots showed high correlation of editing rates of PUM1-APOBEC1 and PUM2-rABE between single labeling and dual labeling cells. (D) Editing rate for PUM1-APOBEC1 and PUM2-rABE in cells. Fewer PUM2 events than PUM1 events are detected, likely due to lower expression level. (E) GSEA analysis of genes with decreased PUM2 binding in PUM1 and PUM2 dual labeling cells.

At least three possibilities could occur between binding sites for PUM1 and PUM2 in cells: independent binding, cooperative binding, or competitive binding. We first checked whether dual expression of both PUM1-APOBEC1 and PUM2-rABE changed binding patterns compared to individual expression. Figure 6B shows an example of PUM1-APOBEC1 and PUM2-rABE editing patterns on *RPL7A*, where we observed similar editing patterns in dual labeling and in single labeling. In another example, *PCNA*, we observed decreased PUM2-rABE editing signal when both PUM1 and PUM2 were overexpressed. Transcriptome-wide, as shown in Figure 6C, editing rates from dual labeling samples were highly correlated with single labeling for both PUM1-APOBEC1 (Pearson’s r = 0.82) and for PUM2-rABE (Pearson’s r = 0.56); only a few hits showed significant differential editing (Figure 6D). In cells expressing both PUM1-APOBEC1 and PUM2-rABE (dual labeling cells), we observed overall slightly higher PUM1-APOBEC1 editing rates than PUM1-APOBEC1 labeling alone, which may be due to slightly higher expression level of PUM1-APOBEC1 in dual labeling cells. We performed GSEA analysis with gene lists ranked by average editing rate changes in dual labeling cells versus cells expressing one deaminase fusion and found that some gene categories had less PUM2 binding in dual labeling cells (Figure 6E), likely due to PUM1 competing for PUM2 binding sites when both proteins were overexpressed. No significant GO enrichment was observed with increased PUM2 binding.

To evaluate whether a single mRNA molecule can be bound by both PUM1 and PUM2, we also performed PacBio long-read sequencing on the same RNA samples used for short-read sequencing. Indeed, we observed that a single RNA molecule can have both A-to-I editing events and C-to-U events, suggesting binding by both PUM1 and PUM2. In the example of the *PCNA* 3’ UTR region in Figure 6B, we examined editing events on the most significant A-to-I site (Position 0) and the most significant C-to-U site (position +30), also a significant C-to-U site further downstream (position +139, not shown in Figure 6B). The editing events observed in these three sites are summarized in Table 1. There was an enrichment of co-editing (1.269 fold) between A-to-I at position 0 and C-to-U at position 139. In contrast, we observed a depletion of co-editing events (0.816 fold) between A-to-I at position 0 and C-to-U at position 30. Taken together, these results showed that by using rABE with a C-to-U RNA deaminase such as APOBEC1, we can probe the binding patterns of two different RNA binding proteins at the same time at the single-molecule level. Moreover, these data indicate that even when one RBP dominates binding, as PUM1 in this case, singlemolecule data can identify RNAs that are selectively bound by PUM2. By combining dual deaminase labeling with long-read sequencing, we identified interactions between RBPs at the single molecule level and across long distances.

**Table 1.**
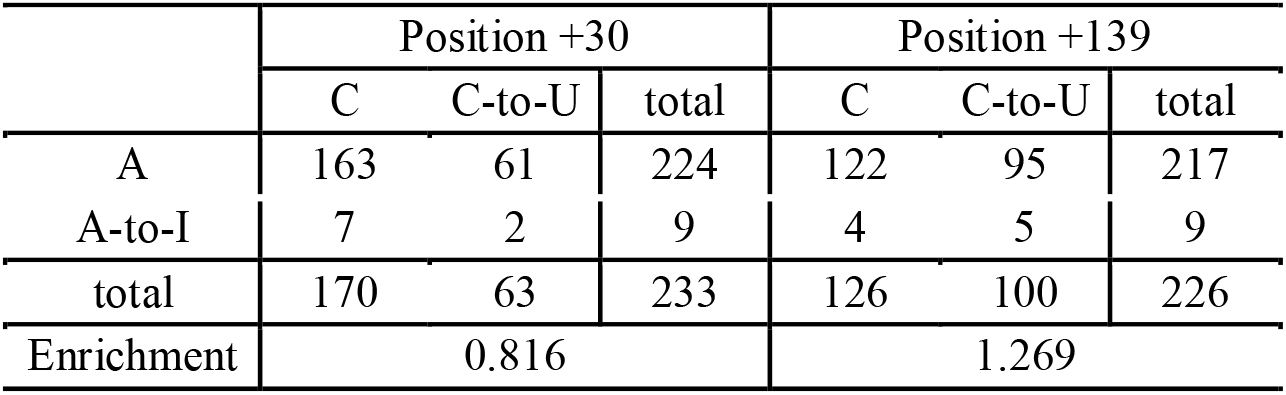
PacBio long-read editing counts at three positions in the PCNA 3’ UTR.

|  | Position +30 |  |  | Position +139 |  |  |
| --- | --- | --- | --- | --- | --- | --- |
|  | C | C-to-U | total | C | C-to-U | total |
| A | 163 | 61 | 224 | 122 | 95 | 217 |
| A-to-I | 7 | 2 | 9 | 4 | 5 | 9 |
| total | 170 | 63 | 233 | 126 | 100 | 226 |
| Enrichment | 0.816 |  |  | 1.269 |  |  |

## Discussion

The systematic understanding of gene expression regulation has been advanced by high-throughput sequencing approaches. For RNA-binding proteins, diverse approaches for capturing RNA-binding sites have been developed for second-generation, short-read sequencing^2–6^. In recent years, new methods are being developed to enable measurement of RNA-binding proteins using third-generation, long-read sequencing, which preserves RNA transcript isoform information and can measure binding at the single-molecule level^44^. In this work, we developed a new adenosine deaminase for RNA molecular recording, an approach we term REMORA (Figure 7).

**Figure 7:**
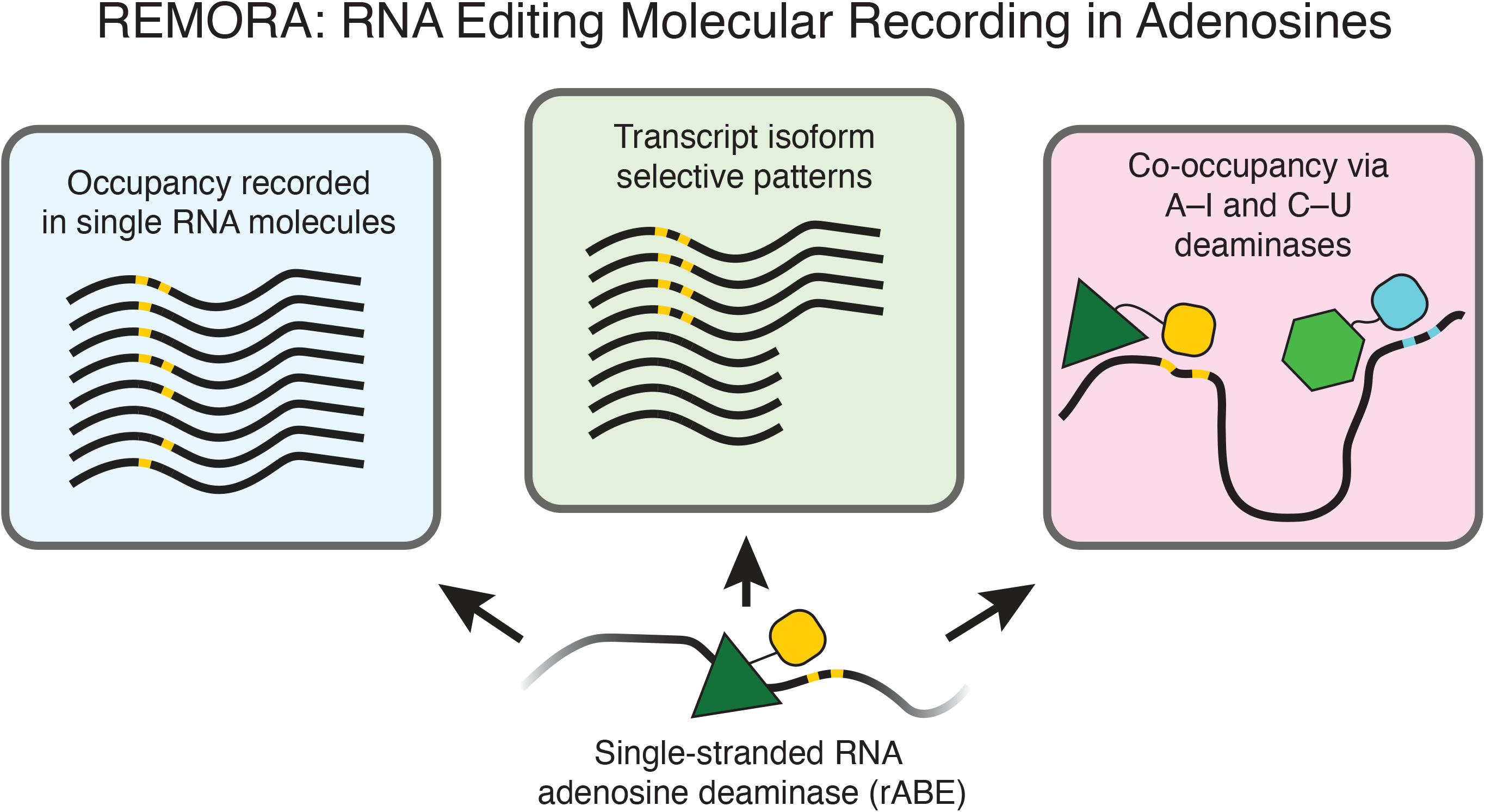
A model of REMORA. Adenosine deamination with rABE enables high fidelity occupancy recording with transcript isoform and single-molecule resolution. When combined with a C–U deaminase such as APOBEC1, co-occupancy and competition between RNA-binding proteins in cells can be measured.

We find that the highly similar Pumilio proteins PUM1 and PUM2 only sometimes compete for the same binding sites in cells. Prior work developed a thermodynamic model of PUM1 and PUM2 binding affinity using high-throughput assessment of PUM1/PUM2 binding affinities in vitro^38^. Jarmoskaite et al. found that PUM1 and PUM2 have indistinguishable sequence preferences *in vitro* and found excellent agreement between their experimentally-derived thermodynamic model and eCLIP-derived PUM1/PUM2 binding patterns. Our finding that PUM1 and PUM2 compete for binding sites in cells is consistent with prior work that concluded PUM1 and PUM2 binding is subsaturating compared to the number of consensus and nonconsensus Pumilio binding elements in the human genome, and therefore occupancy is highly sensitive to changes in PUM levels^38^. Conversely, our data suggesting that competition occurs strongly at some sites and weakly at others may hint at more complexity in in cell binding patterns, perhaps due to the highly skewed abundance distribution of transcripts leading to preferential competition at some sites over others. Interestingly, while there was excellent agreement between thermodynamic modeling and median eCLIP signal in prior work, there was also substantial variance^38^. Continuing to study whether deamination efficiencies by REMORA and related approaches provide a quantitative, direct, comprehensive, and transcript-isoform resolved measurement of RBP occupancy in cells is an exciting direction for the future.

The new RNA deaminase rABE we present in this work originated from directed evolution to develop adenosine base editors for targeting DNA^27^. While ABE7.10 was evolved to target single-stranded DNA, multiple groups observed it has activity against RNA molecules^29–31^. We reasoned that RNA-based directed evolution could improve the activity of ABE7.10 against RNA and devised an RNA-based selection scheme to enrich for more active variants of ABE7.10 against single-stranded RNA. Intriguingly, our selection found that mutation of just one of the ABE7.10 selected variants back to wildtype *E. coli* tadA sequence increased its activity by nearly two-fold. We attempted to discern the mechanism of how this change might increase RNA editing by structural modeling using the *S. aureus* tadA–RNA structure, which suggested possible contacts with the RNA backbone. Future structural work with both rABE and ABE7.10 or related adenosine deaminases bound to both DNA and RNA could identify the origin of nucleic acid selectivity, which could facilitate both development of RNA deaminases and avoiding RNA off-targets in the context of DNA base editing.

The original substrate for ABE7.10 directed evolution was tadA from *E. coli*^27^. Prokaryotic tadA is a homodimeric tRNA deaminase that introduces an inosine into the wobble base of tRNA^Arg-ACG,45^. We explored related tRNA deaminases, including the human ADAT2 and ADAT3 paralogs that install wobble-base inosines in eight distinct tRNAs^46^. We reasoned that since the ADAT2-ADAT3 complex has different tRNA specificity, perhaps it would also function differently as a designed RNA molecular recorder. However, we only detected mild activity with human ADAT2 and ADAT3 (Figure 1). It will be an interesting direction for the future to explore development of human ADAT2-ADAT3 for RNA molecular recording. Moreover, therapeutic introduction of RNA deaminases for programmable deamination might be facilitated using human ADAT2-ADAT3 since human-derived enzymes would be expected to be less immunogenic than those derived from *E. coli* proteins.

Interestingly, the offset in linear sequence space between rABE-induced adenosine deamination and RBP motifs differs in different constructs. In the lambda-BoxB system, dimeric ABE7.10 prefers to deaminate adenosines roughly 11 nt upstream from the BoxB hairpin. When fused to PUM1, rABE also introduced edits at sites about 10-15 upstream from consensus PUM1 motifs. In contrast, Rbfox2-rABE appeared to introduce deamination events both upstream and downstream of consensus Rbfox2 motifs, suggesting that rABE can deaminate adenosines on both sides of Rbfox2 binding sites. An intriguing possibility to investigate in the future is that the structure or dynamics of the RBP that rABE or other deaminase are fused to imparts different distance dependences on deamination, or that RNAs or RBPs can take on multiple conformations might generate conformation-dependent distance dependencies.

Conceptually similar advances to REMORA in DNA have enabled measurement of DNA-related processes at the single-molecule level. DiMeLo-seq uses antibody-tethered enzymes to methylate DNA adjacent to protein binding sites, which revealed single-molecule binding patterns of chromatin-binding proteins and centromerebinding proteins^47^. SAMOSA uses adenine methyltransferases to footprint the location of nucleosomes in single DNA molecules, which identified irregularity in oligonucleosome patterns^48^. As ABE7.10 also functions as an adenine base editor on DNA, it is possible that it could be applied in concert with these approaches for multi-layered encoding. Together, these approaches enable encoding of functional nucleic acid information in third-generation, long-read sequencing with its numerous advantages in long distance information, mapping repetitive regions, disambiguating similar nucleic acid sequences, and single-molecule information.

Expression of active RNA molecular recorders such as rABE or APOBEC1 for extended periods of time is a limitation with the present implementation of REMORA. Long duration expression of active deaminases effectively integrates binding events across the experimental timecourse. As such, binding data inferred from deamination events in this work should be interpreted as binding that was recorded at some point during the duration of the experiment (typically 48 hours post dox induction). Future work in synthetic design of acutely inducible deaminase systems could enable RNA molecular recording with finer time resolution. Alternatively, studying biological processes that can be frozen in time through chemical inhibition or other means could enable higher molecular resolution.

We anticipate that REMORA is compatible with a variety of other information-encoding strategies in RNA. Since rABE is an adenosine deaminase, REMORA is orthogonal to 4-thiouridine (4sU) or 5-ethynyl uridine (5-EU) incorporation and thus can be combined with diverse metabolic labeling approaches^49,50^. Moreover, subcellular localization can be encoded through biotinylation of RNA via APEX-seq^51^, which if combined with REMORA may enable subcellular decoding of RNA binding sites. REMORA combined with fractionated polysome profiling would enable measuring RNA-binding protein association as a function of translation level^7,52,53^. Lastly, as rABE is genetically encodable, cell-type specific RNA-binding protein information can be measured through lineage-specific promoters or other strategies to regulate rABE expression. Long-read compatible, genetically encodable information encoding strategies such as REMORA set the stage for an expanded understanding of singlemolecule RNA biology with cell type resolution as well as manipulation of gene expression through targeted RNA deamination.

## Reagent and data access

Plasmids encoding rABE are available through Addgene (#191383-#191386). Sequencing data will be available through GEO or by request. Analysis code will be posted to Github or is available by request.

## Author Contributions

Conceptualization, Y.L. and S.N.F.; Investigation, Y.L., S.K. B.Q.T., S.N.F., and Y.A.; Writing – Original Draft, Y.L. and S.N.F.; Writing – Review & Editing, Y.L. and S.N.F.; Methodology, Y.L. and S.N.F.; Resources, M.S.O. and K.W.; Formal analysis, Y.L.; Software, Y.L.; Funding Acquisition, S.N.F.; Supervision, S.N.F.

## Acknowledgements

We thank members of the Floor lab for feedback on this work and their continued support. Computation was supported by the UCSF Wynton high-performance computing infrastructure, PacBio sequencing was supported through instrumentation shared with V. Ramani (Gladstone Institutes), high-throughput shortread sequencing was supported by the UCSF Center for Advanced Technology, and flow cytometry was supported by the Parnassus Flow Cytometry Core at UCSF. This work was supported by the UCSF Program for Breakthrough Biomedical Research, funded in part by the Sandler Foundation (to SNF) and the National Institutes of Health DP2GM132932 (to SNF). SNF is a Pew Scholar in the Biomedical Sciences, supported by The Pew Charitable Trusts.

