## Supplemental Figures for "Single molecule co-occupancy of RNA-binding proteins with an evolved RNA deaminase"

### Supplementary Figure 1

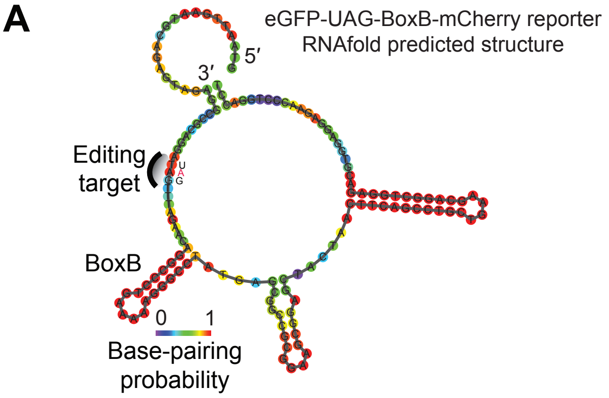


**Supplementary Figure 1.** RNA secondary structure modeling around UAG-BoxB cassette with RNAfold. BoxB forms a stable stem-loop structure, while the UAG stop codon was predicted to be unstructured.

### Supplementary Figure 2

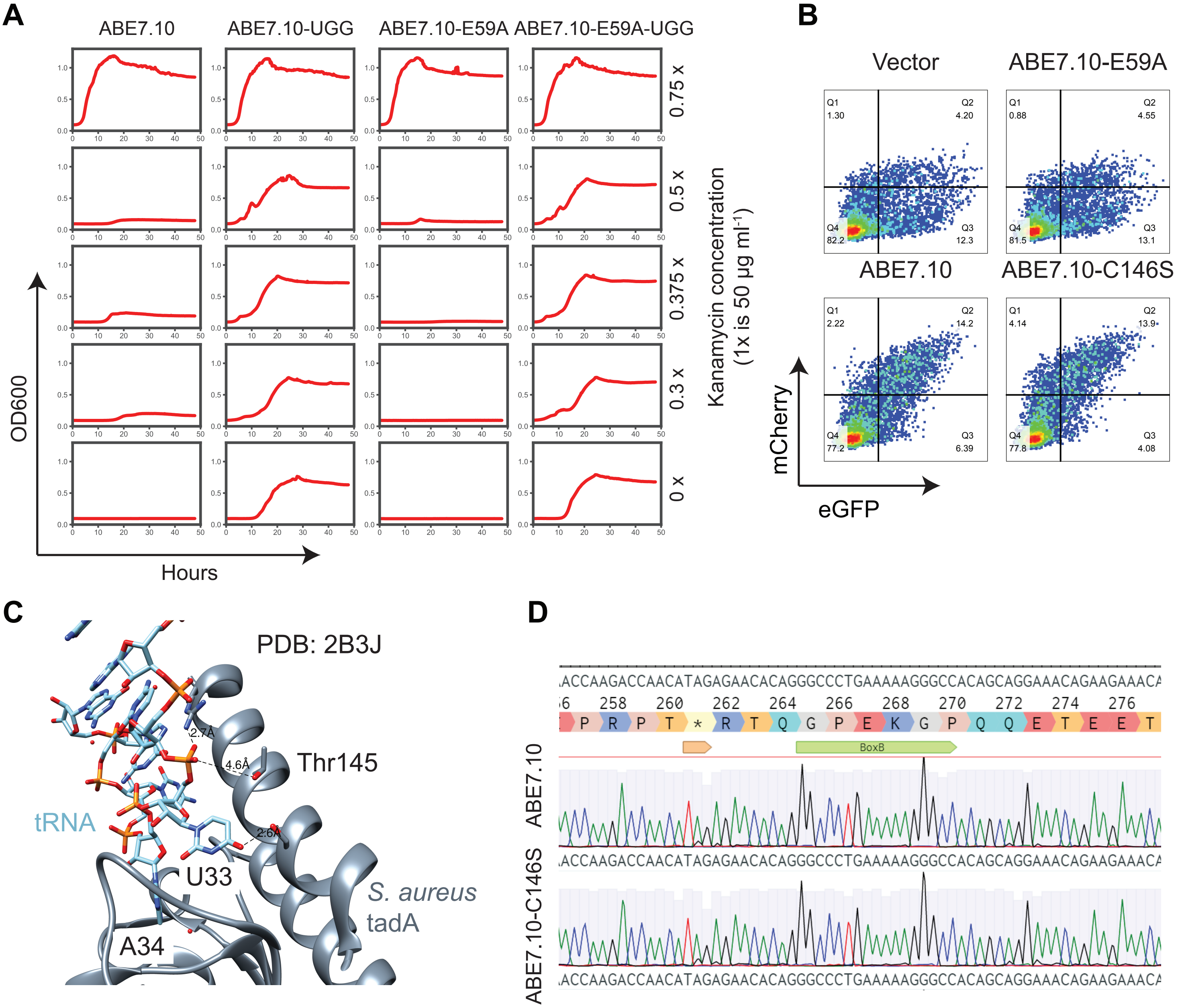


**Supplementary Figure 2.** Directed evolution for an improved ssRNA A-to-I editor. (A) OD600 measurement of E. coli growth curve after transformation of ABE7.10 under different kanamycin concentrations. 0.375 x concentration (18.75 mg/L) was the optimal concentration that allowed growth of ABE.10 expressing cells and inhibited growth of ABE7.10-E59A negative control cells. (B) Scatter plots of flow cytometry measurement of the editing efficiency of ABE7.10 and ABE7.10-C146S. (C) Structural model of the S. aureus tadA protein near Thr145 (corresponding to E. coli Ser146). (D) Sanger sequencing measurement of A-to-I editing rates introduced by ABE7.10 and ABE7.10-C146S on an adenosine-rich UAG-BoxB reporter. Several A-to-I editing were observed flanking both sides of BoxB in different sequence contexts.

### Supplementary Figure 3

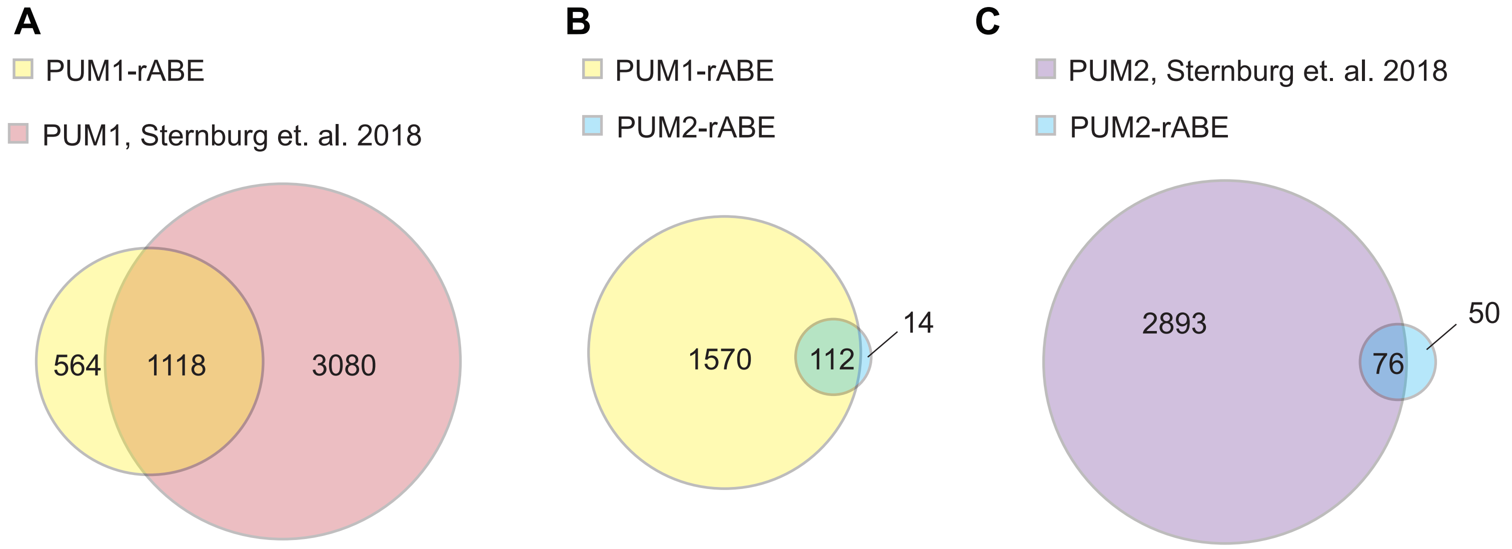


**Supplementary Figure 3.** Venn diagram summary of overlapping gene targets of PUM proteins. (A) Comparison of PUM1-rABE targets (this study) and PUM1 targets identified by CLIP-seq (Sternburg *et al.*[37]). (B) Comparison of PUM1-rABE targets and PUM2-rABE targets. (C) Comparison of PUM2-rABE targets (this study) and PUM2 targets identified by CLIP-seq (Sternburg *et al.*[37]).

### Supplementary Figure 4

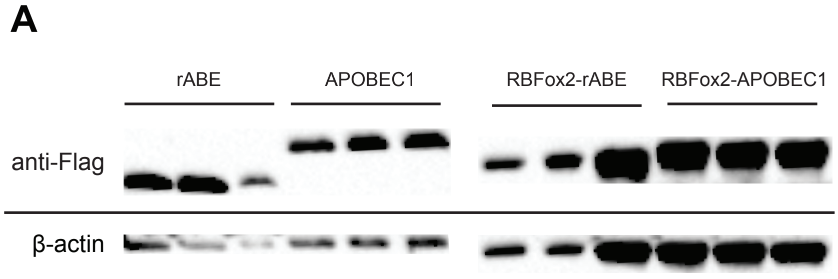


**Supplementary Figure 4.** Western blot measurement of rABE, APOBEC1, Rbfox2-rABE, Rbfox2-APOBEC1 expression levels under 1000 ng/mL doxycycline condition, 3 replicates each. All proteins contain a Flag tag at their C-terminus and are blotted with anti-Flag-HRP.
